# PKCδ mediates high-fat diet-induced increased tonic GABA_A_ receptor current in cardiac vagal motor neurons in the DMV

**DOI:** 10.64898/2026.06.30.735709

**Authors:** Yoko Brigitte Wang, Victor Q. Chen, Matthew McDonald, Christopher D. Romero, Maira Jalil, John N. Campbell, Carie R. Boychuk

## Abstract

Consumption of high fat diet (HFD) is linked to reduced cardiac vagal motor output, a main contributor to progression of cardiovascular disease. HFD for 15 days increases extrasynaptic or tonic gamma aminobutyric acid (GABA) current in cardiac projecting neurons in the dorsal motor nucleus of the vagus (CVN^DMV^), which contributes to dampening cardiac parasympathetic output. However, the mechanism underlying this increased inhibition is unknown. Here, we hypothesize that increased activity of protein kinase C δ isoform (PKCδ) enhances tonic GABA current in CVN^DMV^ after HFD. Whole-cell patch-clamp recording of retrogradely labeled CVN^DMV^ demonstrated that pan inhibition of PKC activity with GFX, and isoform specific inhibition of PKCδ with rottlerin normalize 15-day HFD-induced increases in tonic GABA current, suggesting that PKCδ mediates enhanced tonic inhibition. This effect persisted in the presence of dynasore, a clathrin-mediated endocytosis blocker, indicating that the normalization effect of PKCδ inhibition on tonic current in HFD is likely independent of clathrin-mediated endocytosis. Furthermore, no differences in PKCδ mRNA or protein expression were observed between NFD and HFD, suggesting a post-translational mechanism underpinning increased tonic GABA current after 15 days of HFD. Altogether, this study provides evidence that HFD-induces increased PKCδ activity, but not expression, leading to increased tonic GABAergic inhibition in CVN^DMV^. This increase PKCδ activity could explain the cardiac vagal motor output dampening in CVD and be developed into treatments targeting PKCδ for CVD.

## Introduction

Consumption of a diet high in saturated fat is a significant risk factor for the development of cardiovascular disease (CVD) (Griffin & Lovegrove, 2025; Sacks et al., 2017), including heart failure and hypertension, which all present with reduced cardiac parasympathetic (or vagal) motor tone (Dunlap et al., 2003; Olshansky et al., 2008). Despite this strong correlation between diet high in saturated fat and CVD, understanding of the disease progression is limited, partly because we lack mechanistic models to test in human patients. Studies using animal models, however, confirm that long-term consumption of high-fat diet (HFD) decreases cardiac vagal tone, leading to tachycardia (Bruder-Nascimento et al., 2017; Contreras & Williams, 1989). Emerging evidence further suggests that this HFD-induced dampening of cardiac vagal output occurs as early as 15 days into a HFD regimen, manifesting as resting tachycardia and blunted reflex responsivity (Strain et al., 2023). This early-onset suppression represents a critical window during which vagal dysfunction may precede or drive CVD progression (Carnevali et al., 2013; Costa et al., 2025), making it a particularly important target for mechanistic investigation. Despite this, the complete underlying circuit mechanisms of this HFD-induced blunting of cardiac vagal output remain unknown.

The impact of HFD on cardiac vagal regulation is known to be of central origin (Strain et al., 2023), which sets cardiac vagal motor neurons (CVNs) in the brainstem as a primary target of investigation. CVNs generate cardiac vagal motor output to regulate resting heart rate (HR) and mediate reflex bradycardia. They are located in two brainstem nuclei, nucleus ambiguus and dorsal motor nucleus of the vagus (DMV). Both populations contribute to tonic and reflexive cardiac vagal motor output (Wang, Saunders, et al., 2025). Unlike CVNs in the nucleus ambiguus that produce respiratory-dependent burst firing (Mendelowitz, 1996), CVNs in the DMV (CVN^DMV^) are spontaneously active and exhibit pacemaker-like activity (Strain et al., 2024). CVN^DMV^ reside in the dorsal vagal complex, a major hub that processes and integrates incoming metabolic signals. Given their location and electrophysiological properties, CVN^DMV^ likely serve as signal integrators for metabolic cues, making them susceptible to metabolic perturbations including HFD.

Metabolic challenges disrupt inhibitory neurotransmission in several central nervous system regions (Sandoval-Salazar et al., 2016; Seabrook et al., 2023), including vagal regulatory circuits (Boychuk and Wang, 2026). It is now established that the early impact of HFD includes increased tonic gamma-aminobutyric acid (GABA) currents in CVN^DMV^ (Wang, Dow, et al., 2025). Tonic GABA currents regulate neuronal excitability by producing persistent inhibitory conductance (Semyanov et al., 2004). This persistent current is mediated by GABA type A receptors (GABA_A_Rs) containing δ subunit (GABA_A_Rδ) which are predominantly located at extrasynaptic sites (Brickley & Mody, 2012). Such increases in tonic GABA current lead to reduced neuronal activity, suggesting that elevated tonic GABA_A_Rδ current in CVN^DMV^ may contribute to decreased cardiac parasympathetic output after 15-days of HFD. Importantly, deletion of GABA_A_Rδ in vagal motor neurons eliminates the impact of early HFD on cardiac vagal motor output (Strain et al., 2024), establishing GABA_A_Rδ as a critical mediator of HFD-induced increases in tonic GABA current.

Protein kinase C (PKC) is a serine/threonine kinase that regulates GABA_A_Rs phosphorylation (Jacob et al., 2008), which can alter GABA_A_R functional activity through changes in membrane stability and expression, receptor internalization, and receptor kinetics (Jacob et al., 2008; Nakamura et al., 2015). Critically, disruption of GABA_A_R phosphorylation contributes to disease pathophysiology (Garcia et al., 2023; Vien et al., 2015). PKC modulates extrasynaptic GABA_A_Rs and tonic inhibition in a neuronal cell-type dependent manner (Abramian et al., 2010; Brandon et al., 2000; Bright & Smart, 2013). In DMV neurons, GABA_A_Rδ requires PKC activity to facilitate state-dependent effects on tonic GABA current (Littlejohn & Boychuk, 2021), suggesting that PKC is involved in regulating tonic inhibition in this neuronal population at least under certain conditions.

PKC, itself, has three main isoform classes, conventional (α, βI, βII, and γ), novel (δ, ε, η, and θ) and atypical (ζ, ι, and λ) (Mochly-Rosen et al., 2012). Cholinergic DMV neurons, which include CVNs, express transcripts of PKC α, β, δ, ε, ζ, and ι isoforms (Tao et al., 2021). Among these PKC isoforms, PKCδ expression is upregulated after HFD (Bezy et al., 2011), and its upregulation can promote increased in tonic GABA current in disease states at least in other brain regions (Choi et al., 2008). Therefore, we hypothesized that HFD increases PKCδ activity in CVN^DMV^, promoting membrane stabilization of GABA_A_Rδ, leading to enhanced tonic GABA current.

Here, we used ex vivo whole cell patch-clamp electrophysiology, immunohistochemistry and endocytosis assays to investigate the role of PKCδ in driving increased tonic GABA current in CVN^DMV^ after HFD. We found that inhibition of PKCδ normalized HFD-induced tonic GABA current in CVN^DMV^. This effect is independent of clathrin-dependent endocytosis, implicating either reduced exocytosis or receptor kinetics as possible mechanisms of PKCδ enhancement of GABA_A_Rδ function. PKCδ mRNA and protein expression levels was also unchanged, suggesting that PKCδ is post-translationally modified to regulate GABA_A_R functional activity. Together, our findings demonstrate that PKCδ mediates the increase in tonic GABA current in CVN^DMV^ after 15-days of HFD, providing a potential cellular mechanism underlying early cardiac parasympathetic dysfunction induced by HFD.

## Results

### PKCδ mediates HFD-induced tonic GABA current increase in CVN^DMV^

In DMV neurons, PKC is important for GABA_A_Rδ function (Littlejohn & Boychuk, 2021) although whether this applies specifically to CVN^DMV^ is unknown. Identifying whether PKC-dependent activity drives tonic GABA currents in CVN^DMV^ would provide mechanistic insights into HFD-induced tonic GABA currents increases and cardiac parasympathetic dysfunction. Since PKCδ can facilitate the impact of HFD in other tissue types, initial investigations confirmed the presence of PKCδ mRNA in CVN^DMV^ under NFD conditions using RNAscope (Figure 1A) and single-cell RT-PCR (Figure 1B). Immunohistochemistry further confirmed the expression of PKCδ at the protein level (Figure 1C), demonstrating that CVN^DMV^ express PKCδ at both transcript and protein levels.

**Figure 1.**
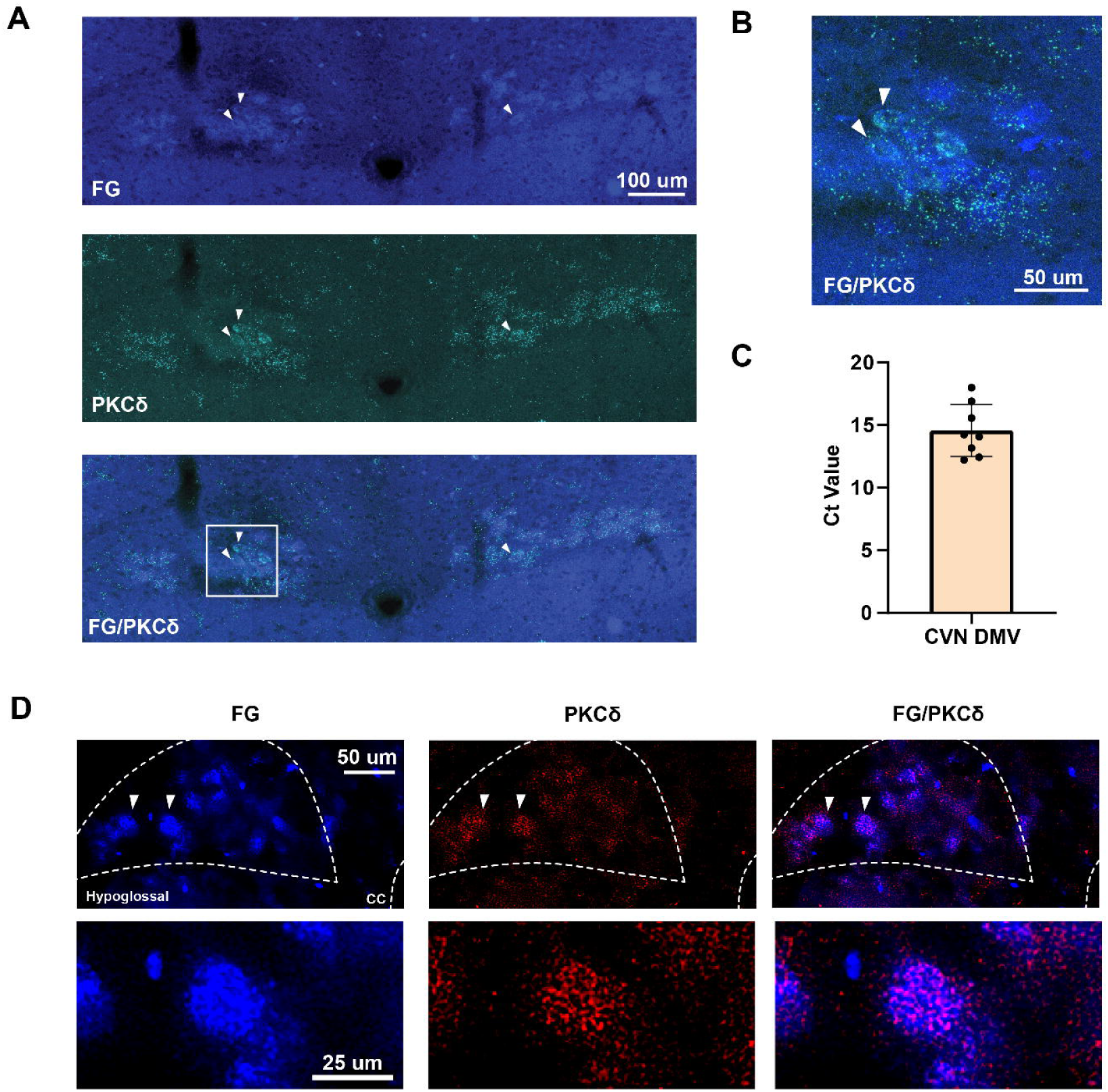
Expression of PKCδ mRNA and protein in CVN^DMV^ in NFD. A) Representative images of PKCδ mRNA expression in CVN^DMV^ (traced with fluorogold) of NFD mice (n=2 mice), B) Enlarged representative image of PKCδ mRNA expression in CVN^DMV^ of NFD mice, C) Single-cell RT-PCR Ct value of PKCδ mRNA in patched CVN^DMV^ (n=8 cells), D) Representative images of PKCδ protein expression in CVN^DMV^ of NFD mice (n=4 mice), bottom row: close up image of PKCδ protein expression in CVN^DMV^. FG, fluorogold; PKCδ, protein kinase C δ; CVN, cardiac vagal motor neurons; DMV, dorsal motor nucleus of the vagus; CC, central canal.

To determine whether PKC could be responsible for the functional impact of HFD on CVN output, retrograde cardiac tracing (Fig 2A, B) was paired with ex vivo whole cell patch-clamp recordings in brainstem slice preparation under voltage-clamp configuration to determine if PKC, and its specific isoform, PKCδ, are involved in GABAergic currents in CVN^DMV^. Similar to previous report (Wang, Dow, et al., 2025), 15 days of HFD significantly increased tonic GABA current in CVN^DMV^ (Figure 2D, NFD_control vs_ HFD_control_ = 0.98±0.27 vs 2.01±0.29 pA/pF, p=0.007, n=11, 8). Application of GF 109203X (GFX; 500nM), a pan-PKC inhibitor (Toullec et al., 1991), in the perfusate abolished the increase in tonic GABA current density in CVN^DMV^ induced by 15 days of HFD (HFD_GFX vs control_ = 0.85±0.30 vs 2.01±0.29 pA/pF, p=0.006, n=7, 8, Figure 2D), restoring tonic current to similar density as in CVN^DMV^ from NFD mice (vs NFD_control_ = 0.98±0.27 pA/pF, p=0.72, n=11; vs NFD_GFX_ = 0.79±0.15 pA/pF, p=0.88, n=8). Pan-inhibition of PKC had no effect on tonic current in CVN^DMV^ from NFD (Figure 2D), similar to the lack of effects at baseline in unlabeled DMV neurons (Littlejohn & Boychuk, 2021).

**Figure 2.**
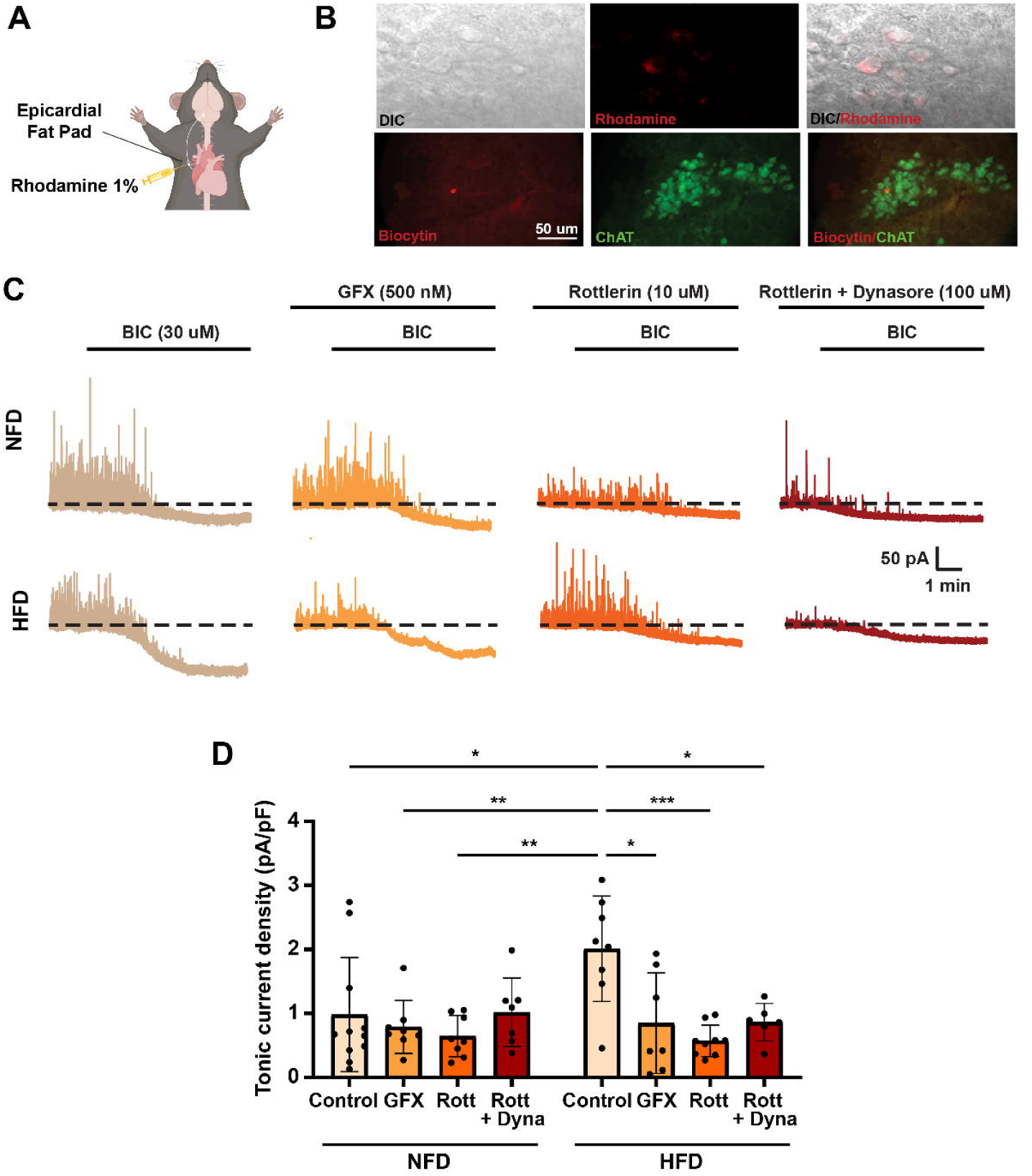
Inhibition of PKCδ normalized HFD-induced increased of tonic current in CVN^DMV^. A) Illustration of retrograde cardiac injection using rhodamine 1% to label CVN^DMV^, B) Top row, representative image of rhodamine-labelled CVN^DMV^ for patch-clamp electrophysiology. Bottom row, post-recording confirmation of biocytin-filled neurons and ChAT, C) Representative traces of tonic GABA current in NFD and HFD groups under control condition and different drugs, D) Tonic current density of CVN^DMV^ in NFD and HFD groups under control condition (NFD n=11 cells, HFD n=8 cells), GFX (NFD n=8 cells, HFD n=7 cells), rottlerin (NFD n=8 cells, HFD n=8 cells), rottlerin + dyansore (NFD n=7 cells, HFD n=6 cells), two-way ANOVA (p_diet_ = ns, p_drugs_ = 0.022, p_diet x drugs_ = 0.0005) with Sidak’s multiple comparisons test. * p≤0.05, ** p≤0.01, *** p≤0.001. Data is presented as mean ± SD. DIC, differential interference contrast; ChAT, choline acetyltransferase; NFD, normal fat diet; HFD, high fat diet; BIC, bicuculine methiodide; GFX, GF 109203X, Rott, rottlerin; Dyna, dynasore.

To test then whether PKCδ specifically mediates the HFD-induced increase in tonic GABA current in CVN^DMV^, rottlerin, a PKCδ inhibitor (10 µM) (Kuver et al., 2012), was applied in the perfusate. Tonic current density did not differ between NFD groups with or without rottlerin. However, PKCδ inhibition normalized tonic current density in CVN^DMV^ from HFD mice (HFD_Rottlerin vs control_ = 0.57±0.08 vs 2.01±0.29 pA/pF; p<0.0001; n=9, 8; Figure 3B) to levels similar to those observed in CVN^DMV^ from NFD mice (NFD_Control_ = 0.98±0.27 pA/pF, p=0.18, n=11; NFD_Rottlerin_ = 0.65±0.11 pA/pF; p=0.28, n=9). These findings suggest that elevated PKCδ activity mediates HFD-induced increases in tonic GABA current in CVN^DMV^.

**Figure 3.**
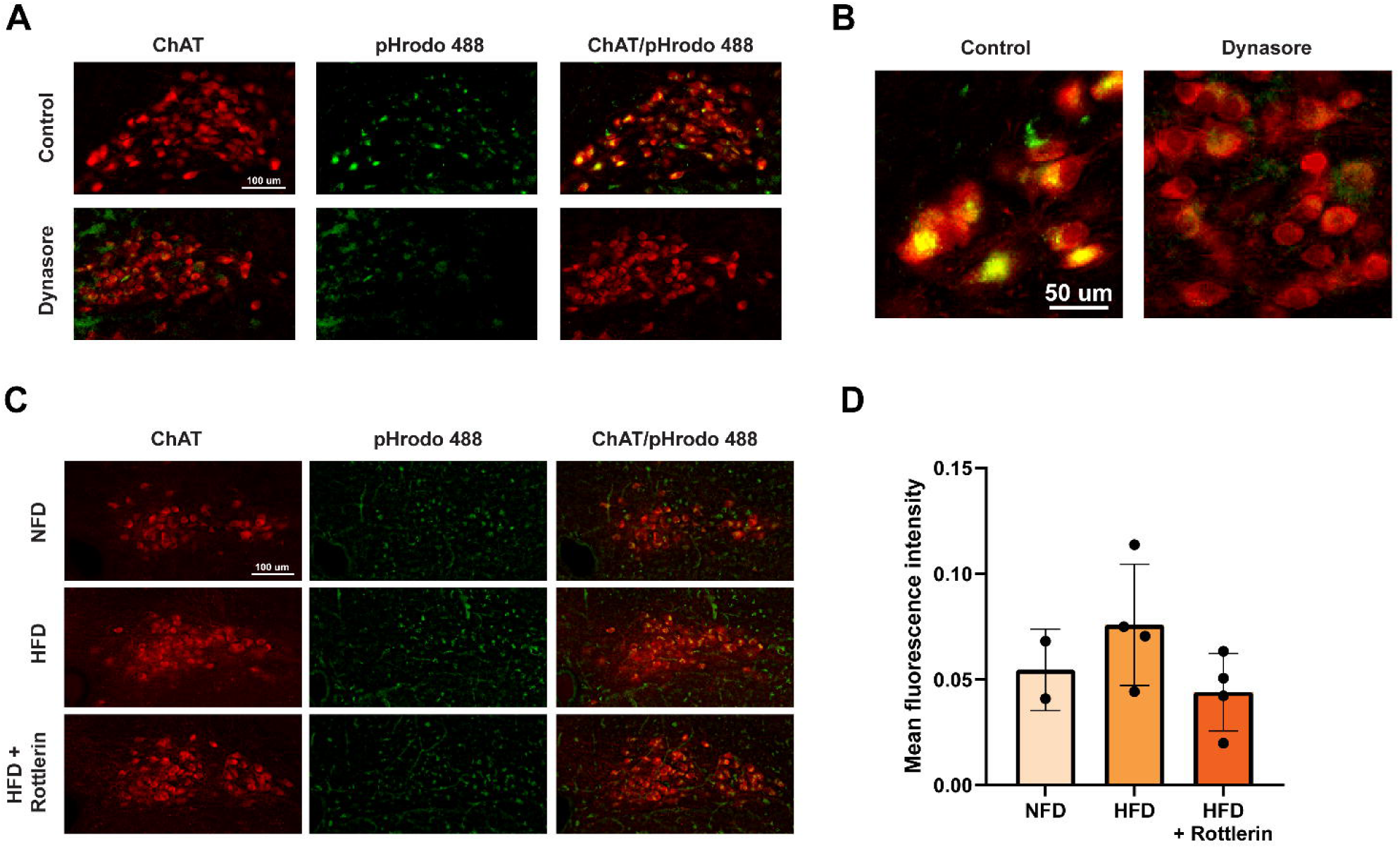
HFD does not alter global endocytosis in ChAT-positive neurons in the DMV. Representative images of pHrodo signals in ChAT-positive neurons in the DMV in A) control and dynasore groups, B) close up image of control and dynasore groups, C) NFD, HFD and HFD+Rottlerin groups, D) No significant difference in mean fluorescence intensity of pHrodo between NFD (n=2 mice), HFD (n=4 mice), and HFD + Rottlerin (n=4 mice) groups. Data is presented as mean ± SD. ChAT, choline acetyltransferase; NFD, normal fat diet; HFD, high fat diet.

### Reduction of tonic current after PKC**δ** inhibition in HFD is independent of clathrin-mediated endocytosis

Phosphorylation of GABA_A_Rs can reduce receptor internalization, promote insertion, and change receptor kinetics dependent on subunits being phosphorylated (Abramian et al., 2014; Abramian et al., 2010; Jacob et al., 2008). It is possible then that increased PKCδ activity drives GABA_A_R phosphorylation to prevent receptor internalization from the membrane. As a consequence, this could increase the number of receptors present on the membrane, leading to enhanced tonic GABA current. Therefore, whether the reduction in tonic current following PKCδ inhibition is due to receptor internalization from the membrane via endocytosis was examined.

GABA_A_Rs are recycled from the membrane primary via clathrin-dependent endocytosis (Jacob et al., 2008). Here, an endocytosis assay in brainstem slice using pHrhodo conjugated dextran was performed to confirm the usefulness of this technique in DMV neurons (Fig 3A). Assay validation was done by adding dynasore (80 µM), a dynamin GTPase inhibitor that prevents clathrin-dependent endocytosis, to the perfusate (Hofmann & Andresen, 2017; Macia et al., 2006). pHrodo intensity density was reduced after dynasore application suggesting that pHrodo was being endocytosed via a clathrin-dependent mechanism (Figure 3A, B). This assay was then used to test the impact of HFD on endocytosis. There was no difference in pHrodo uptake as measured by mean intensity between NFD, HFD and HFD with rottlerin (Figure 3C, D), indicating that HFD and PKCδ inhibition in HFD does not influence global endocytosis in cholinergic DMV neurons.

To investigate whether blocking endocytosis specifically in CVN^DMV^ diminishes the effect of PKCδ inhibition on tonic GABA current of HFD mice, patch clamp electrophysiology was again performed. This time dynasore (100µM) was applied during whole cell patch clamp experiments through addition to the perfusate together with rottlerin. Blockade of clathrin-dependent endocytosis did not abolish the normalization effect of PKCδ inhibition on tonic GABA current in CVN^DMV^, and there was no difference between NFD and HFD group (Figure 2D). However, since clathrin-dependent endocytosis is also critical for pre-synaptic terminal release (Milosevic, 2018; Royle & Lagnado, 2010), the frequency of IPSCs was also significantly reduced compared to control in each diet group (NFD = 0.53±0.78 vs 3.16±3.14 Hz, p = 0.046, n= 7 cells; HFD = 3.97±2.15 vs 0.48±0.51 Hz, p = 0.003, n=6 cells; Kruskal-Wallis with Dunn’s multiple comparisons test).

### PKCδ inhibition decreases IPSC decay time in CVN^DMV^ of NFD

It is established that 15-day of HFD does not alter IPSC frequency, amplitude or decay time in CVN^DMV^ (Wang, Dow, et al., 2025). Consistent with this, the present study confirms that 15-day HFD did not change phasic current frequency, amplitude and decay time in CVN^DMV^ compared to CVN^DMV^ from NFD mice in control conditions without pharmacological interventions (Figure 4B, 4E, 4F). To determine whether PKCδ inhibition impacts phasic GABAergic neurotransmission, IPSC frequency, amplitude and decay time were then measured in CVN^DMV^ from NFD and HFD groups. No difference in IPSC frequency existed after application of either GFX or rottlerin, indicating that presynaptic input to CVN^DMV^ remains likely unaffected by PKC activity (Figure 4B). GFX did not alter IPSC amplitude (Figure 4D) or decay time (Figure 4F) in both CVN^DMV^ from NFD and HFD groups, suggesting that pan-inhibition of PKC does not induce postsynaptic modification in CVN^DMV^. Rottlerin did not alter IPSC amplitude in NFD and HFD groups.

**Figure 4.**
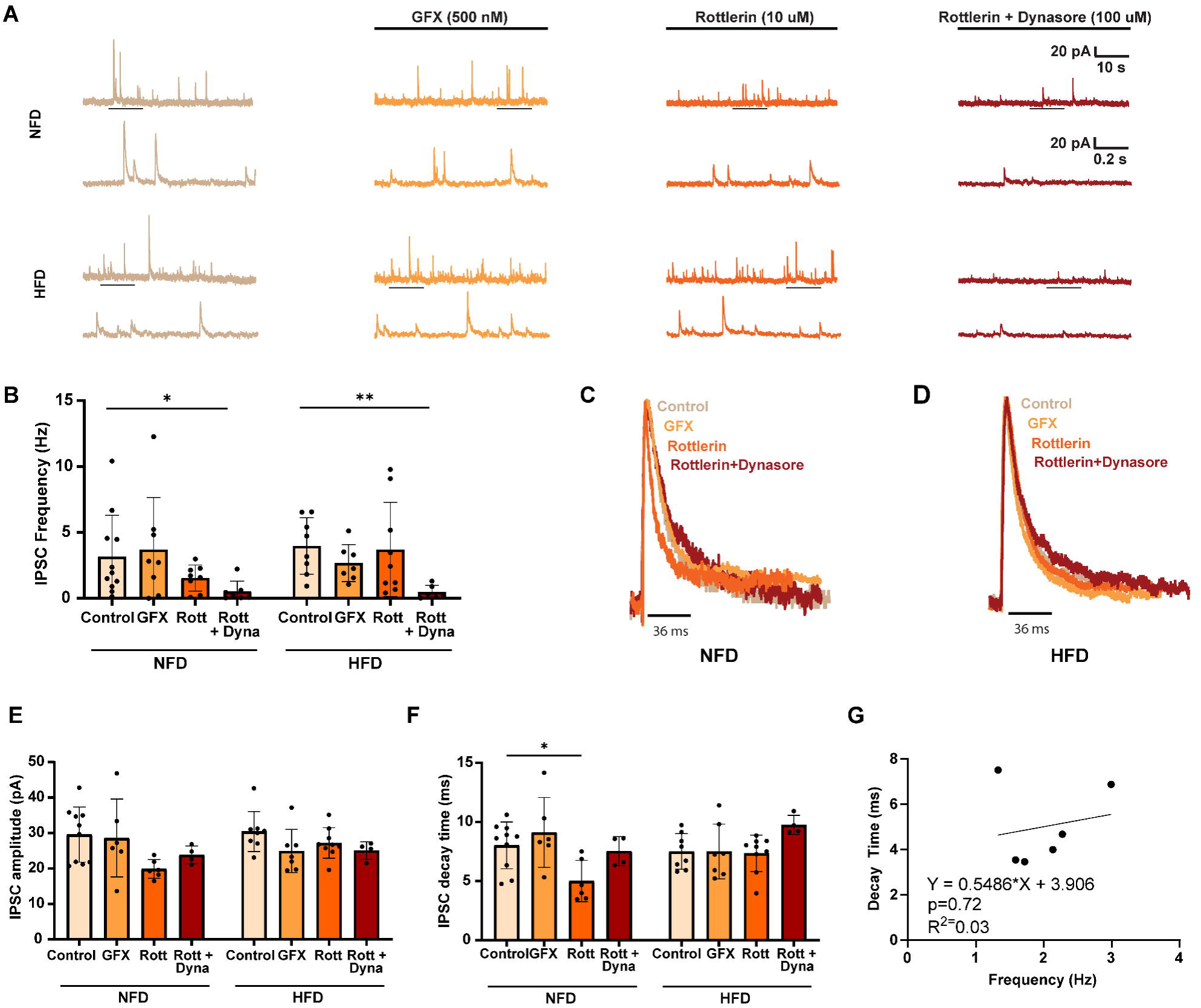
Inhibition of PKCδ decreases decay time in CVN^DMV^ of NFD group. A) Representative traces of IPSC GABA current in CVN^DMV^ under control conditions and different drugs, B) IPSC frequency of CVN^DMV^ in control (NFD n=11, HFD n=8), GFX (NFD n=8, HFD n=7), rottlerin (NFD n=8, HFD n=8), rottlerin + dynasore (p=0.046, NFD n=7, p=0.003, HFD n=6; Kruskal-Wallis with Dunn’s multiple comparisons test), C) Representative traces of a single IPSC event from NFD under control (n=10), GFX (n=6), rottlerin (n=6), and rottlerin+dynasore (n=4), D) Representative traces of a single IPSC event from HFD under control (n=8), GFX (n=7), rottlerin (n=8) and rottlerin+dynasore (n=4), E) IPSC amplitude, F) IPSC decay time (p=0.038, Kruskal-Wallis with Dunn’s multiple comparisons test), G) linear regression of frequency and decay time in NFD_Rottlerin_ group. Data is presented as mean ± SD. * p≤0.05, ** p≤0.01. NFD, normal fat diet; HFD, high fat diet; BIC, bicuculine methiodide; GFX, GF 109203X, Rott, rottlerin; Dyna, dynasore.

However, rottlerin application significantly reduced decay time in CVN^DMV^ from NFD mice when compared to CVN^DMV^ from NFD mice under control aCSF conditions (5.01±1.76 vs 8.02±1.98 ms, p=0.038, control = 10 cells, rottlerin =6 cells, Kruskal-Wallis with Dunn’s multiple comparisons test, Figure 4F). A linear regression confirmed no relationship existed between the reduction in IPSC frequency and decay time (R^2^=0.03; p=0.72; Figure 4G), ensuring that reduced IPSC frequency did not impact the impact of rottlerin. No effect on decay time was observed in HFD.

### Increased of PKC**δ** activity in CVN^DMV^ after HFD is independent of transcription and translation

HFD can increase PKCδ mRNA and protein expression (Bezy et al., 2011). Single-cell RT-PCR results from CVN^DMV^ showed that 15-days of HFD did not alter mRNA levels compared to CVN^DMV^ from NFD mice (NFD vs HFD = 1.04±0.63 vs 0.51±0.25, NFD n=7 cells, HFD n=8 cells, Figure 5B). No difference in PKCδ protein expression in CVN^DMV^ was observed between diet groups using immunofluorescence analysis (NFD vs HFD = 7102±4483 vs 10614±4376 IU/px^2^, NFD n=4 mice, HFD n=3 mice, Figure 5D). Taken together, these suggest that the mechanisms underlying PKCδ activity in HFD are likely from post-translational modifications.

**Figure 5.**
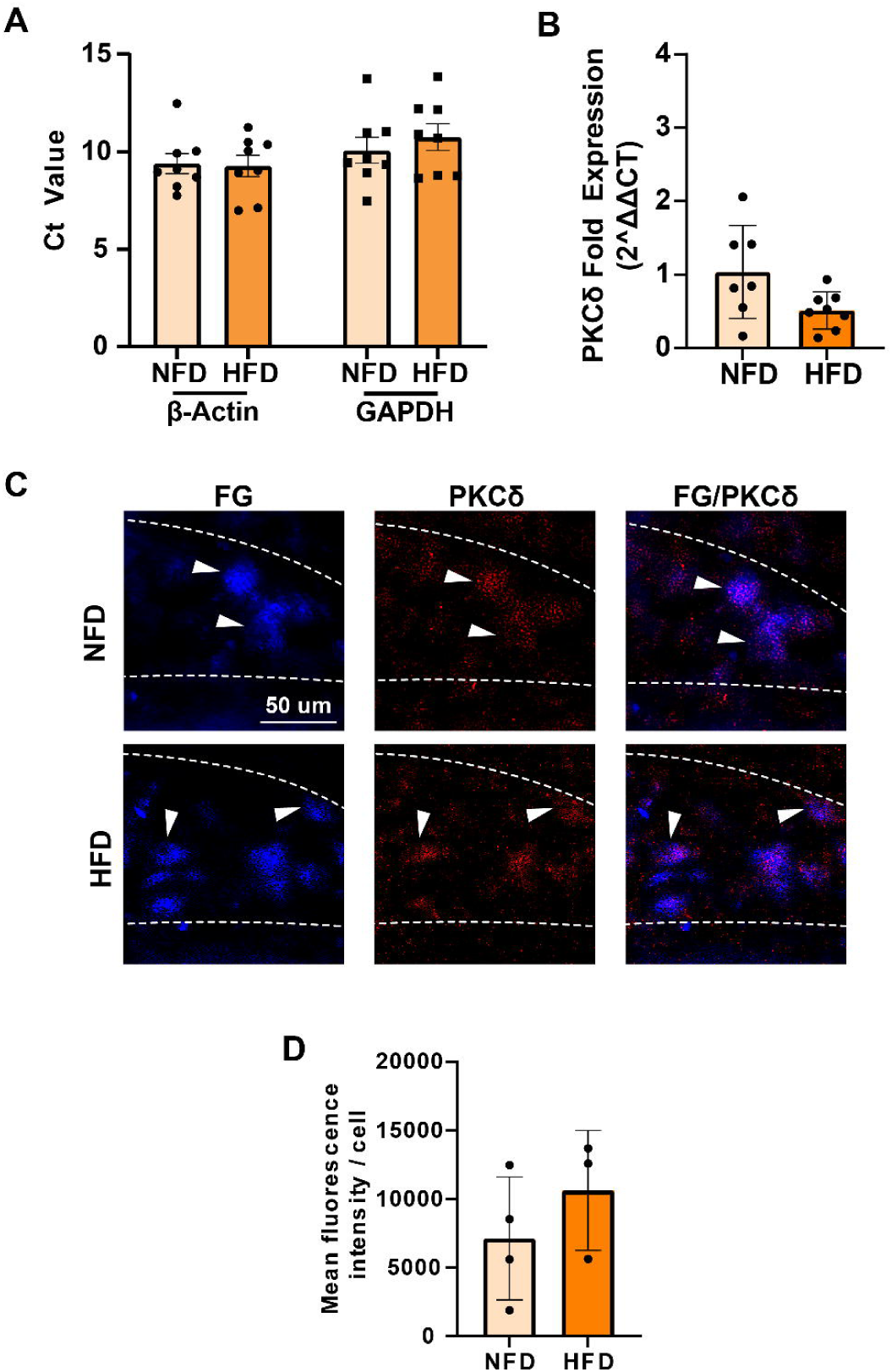
No difference in PKCδ mRNA and protein expression between NFD and HFD groups. A) Single-cell qRT-PCR Ct value of housekeeping genes from recorded CVN^DMV^ in NFD (n=7 cells) and HFD (n=8 cells) groups, B) PKCδ mRNA fold-expression of CVN^DMV^ in NFD and HFD, C) Representative images of PKCδ protein expression in CVN^DMV^ labelled with fluorogold, D) Mean fluorescence intensity of PKCδ per CVN^DMV^ in NFD and HFD. Data is presented as mean ± SD. NFD, normal fat diet; HFD, high fat diet, PKCδ, protein kinase C δ.

## Discussion

The present study identifies a critical role of PKCδ in the regulation of tonic GABA currents in CVN^DMV^ and reveals that PKCδ activity is required for the HFD-induced increase in tonic inhibition in this population. This effect is independent of clathrin-mediated endocytosis, suggesting that PKCδ promotes extrasynaptic GABA_A_Rs membrane availability through a mechanism distinct from receptor internalization. Furthermore, HFD does not significantly alter PKCδ mRNA or protein expression levels in CVN^DMV^, which indicates that HFD acts at a post-translational level to modulate PKCδ activity. Altogether, these findings demonstrate a cellular mechanism by which 15 days of HFD disrupts tonic GABA currents in CVN^DMV^, contributing to impaired cardiac parasympathetic output.

### Interaction of PKCδ and extrasynaptic GABA_A_Rs

PKC regulates GABAergic neurotransmission through phosphorylation of both synaptic and extrasynaptic receptors. While PKC-dependent regulation of synaptic GABA_A_Rs is well established (Field et al., 2021), PKC regulation of extrasynaptic GABA_A_Rs, particularly in the brainstem, remains less understood. For example, in hippocampal dentate granule cells, PKC activation increased tonic GABA current (Abramian et al., 2014; Modgil et al., 2019; Parakala et al., 2019), in part through phosphorylation on the α4 (S443) (Abramian et al., 2010) or β3 subunits (Parakala et al., 2019). Similarly, activation of PKC with PMA elevated tonic GABA current in hypothalamic paraventricular neurons (Womack et al., 2007). In contrast, PKC activation dampened tonic current in hippocampal CA1 pyramidal neurons (Rombo et al., 2016), likely through direct phosphorylation of the β2 (S410) subunit (Bright & Smart, 2013). Previous reports also demonstrate that PKC did not change tonic current, or phosphorylation, of extrasynaptic receptors containing α4 and δ subunits in cultured cortical neurons (Bohnsack et al., 2016; Carlson et al., 2016). Beyond differences between neurons themselves, specific PKC isoforms also differentially phosphorylate GABA_A_Rs (Kia et al., 2011), further contributing to divergent modulation. Taken together, these previous reports confirm that PKC-dependent phosphorylation of extrasynaptic GABA_A_Rs produce divergent effects on neuronal populations arising at least in part from significant differences in GABA_A_R subunit stoichiometry and thereby composition and the location of phosphorylation.

Similar to previous reports from unlabeled DMV neurons (Littlejohn & Boychuk, 2021), PKC inhibition with either GFX or rottlerin had no significant effect on tonic current In CVN^DMV^ from NFD mice. In striking contrast, PKC inhibition with either GFX or rottlerin significantly reduced tonic current in CVN^DMV^ from HFD mice. Similar state-dependent actions of PKC activity on GABA_A_R tonic current are not uncommon. Using PKCδ as an example, PKCδ mediates an increase in tonic GABA current during acute ethanol exposure (Choi et al., 2008), but also facilitates the reduction of tonic GABA_A_R current during ethanol withdrawal since its inhibition abolished the ability of neurons to response to withdraw through reduces in tonic GABA_A_R current (Chen et al., 2018). State-dependent inhibitory signaling in DMV is a proposed mechanism of neuronal plasticity in DMV (Browning et al., 2004; Browning & Travagli, 2009; Littlejohn & Boychuk, 2021), but the present study extends our understanding of state-dependent inhibition in the brainstem to include not only an interaction between GABA_A_Rs with diet and metabolic cues, but also PKC isoform specific actions.

An outstanding question remains: which subunits interact with PKCδ in CVN^DMV^? To date, evidence on the interaction of PKC isoforms with specific GABA_A_R subunits is lacking. Among conventional isoforms, PKCβII phosphorylates β1 (S409) and β3 (S408/409) (Brandon et al., 1999). To our knowledge, limited data exists on which subunit PKCδ specifically phosphorylates to achieve its actions. However, using a heterologous expression system, PKCδ reduced surface expression of receptors containing α4β2δ (Kuver et al., 2012), implicating these three subunits. In CVN^DMV^, mRNA expression of α1 and δ subunits are present, but not α4 (Wang, Dow, et al., 2025). Therefore, it is likely that PKCδ acts on GABA_A_R containing α1, β2 and/or δ subunits. Further studies are warranted to determine the specific target subunit of PKCδ in CVN^DMV^.

### PKCδ mechanism of actions driving tonic GABA current in CVN^DMV^

It is well established that phosphorylation of GABA_A_Rs determines receptor function by regulating their membrane surface stability, spatio-temporal localization and channel kinetics. Since surface expression is a proposed mechanism for the impact of HFD on CVND^MV^, the role of receptor internalization through clathrin-mediated endocytosis was explored here. Neither immunohistochemical analysis or electrophysiology results support a PKCδ-dependent increase in clathrin-mediated endocytosis as a mechanism for rotterlin’s ability to abolish the impact of HFD to increase tonic current in CVN^DMV^. It should be noted that this interpretation is complicated by the near-absence of IPSC frequency under dynasore application, suggesting insufficient ambient GABA to induce tonic current (HONG GAO). A follow-up study incorporating GABA into the preparation is therefore required to determine whether PKCδ regulation of tonic current is independent of endocytosis.

In addition then to receptor internalization, GABA_A_R surface expression is determined by two other main factors: receptor insertion and lateral diffusion between synaptic and extrasynaptic sites. These factors determine how many receptors are present on the membrane, how long the receptors stay, and where the receptors are located. All of which are critical for the spatio-temporal regulation of GABA_A_Rs to maintain phasic and tonic inhibitions. Understanding which of these pathways PKCδ acts through is critical to determine how tonic GABA inhibition is regulated in CVN^DMV^.

Phosphorylation can also promote membrane insertion of GABA_A_R, which may increase exocytosis and receptor abundance on the membrane. Evidence suggests that PKC-dependent phosphorylation of α4 (S443) subunit promotes membrane insertion of extrasynaptic GABA_A_Rs (Abramian et al., 2014; Abramian et al., 2010). Although α4βxδ is a common heteropentameric configuration for extrasynaptic GABA_A_Rs, no α4 subunit mRNA expression was detected in CVN^DMV^ (Wang, Dow, et al., 2025). Thus, it is less likely that PKC-dependent phosphorylation of α4 plays a part in PKCδ action in CVN^DMV^ to increase tonic GABA current via receptor insertion in HFD.

Phosphorylation promotes clustering of GABA_A_Rs which strengthens phasic or tonic current (Petrini et al., 2004; Petrini et al., 2014). GABA_A_Rs are inserted into the membrane at the extrasynaptic site and recruited to the synaptic site through lateral diffusion (Bogdanov et al., 2006). This process is largely dependent on the scaffolding proteins, gephyrin and collybistin (Jacob et al., 2008), binding to phosphorylated GABA_A_R α2 (S359) or β3 (S383) subunit (Nakamura et al., 2020; Petrini et al., 2014). Although phosphorylation of those subunits is typically dependent on protein kinase A and CaMKII, respectively (Nakamura et al., 2020; Petrini et al., 2014), as previously mentioned the actions of PKC can be cell type specific.

Given that PKC inhibition with rottlerin significantly reduced the decay time of phasic IPSCs in CVN^DMV^ neurons from NFD mice, but not from HFD mice, it is tempting to speculate that HFD promotes the redistribution of PKCδ-sensitive GABA_A_Rs away from the synapse. Because PKCδ inhibition shortened phasic IPSC decay kinetics, these receptors may normally contribute to prolonged inhibitory signaling, similar to their proposed role in mediating tonic inhibition. However, this hypothesis remains speculative, and additional studies specifically examining GABA_A_R localization and function in CVN^DMV^ neurons will be required to establish this mechanism.

Besides orchestrating their spatio-temporal localization, phosphorylation of specific subunits also determines GABA_A_Rs channel kinetics. Studies have shown that PKC phosphorylates α4 subunit which increases α4-containing GABA_A_Rs stability on the membrane, preventing tonic current rundown (Abramian et al., 2010). However, CVN^DMV^ does not express α4 subunit mRNA (Wang, Dow, et al., 2025) and their GABA_A_Rδ subunit configuration is unknown. CVN^DMV^ expresses α1 subunit, which has been reported to pair with δ to form a α1βxδ tonic receptor in hippocampal interneurons (Glykys et al., 2007). It is likely then that δ subunit pairs with α1 in CVN^DMV^. Regardless, whether PKCδ-dependent phosphorylation alters GABA_A_Rδ channel kinetics and thereby contribute to an increase in tonic current in HFD, remains to be determined.

Beyond receptor level regulation, tonic GABA current also depends on cellular chloride gradient. Potassium chloride cotransporter-2 (KCC2) facilitates chloride (Cl^-^) and potassium (K^+^) ions extrusion from the cell, which maintaining a low concentration of intracellular Cl^-^ that allows Cl^-^ influx upon GABA_A_R opening to produce hyperpolarization. Phosphorylation of KCC2 is regulated by multiple kinases, including PKC (Kahle et al., 2013). PKC directly phosphorylates at KCC2 serine 940 residue increases its surface stability and reduces internalization (Lee et al., 2007). Given that PKCδ is activated under HFD conditions, it is plausible that PKCδ-dependent phosphorylation of KCC2 contributes to enhanced tonic inhibition in CVN^DMV^ by altering chloride homeostasis.

### PKCδ and phasic GABA current in CVN^DMV^

This study showed that inhibition of PKCδ does not alter IPSC frequency and amplitude in NFD and HFD. This indicates that PKCδ activity less likely to contribute in presynaptic GABA release and the abundance of active receptors at the synaptic site. Indeed, a significant reduction was observed on decay time after application of rottlerin in NFD group. However, this was only significant compared to the effect of pan-PKC inhibition in NFD and when endocytosis and PKCδ were blocked in HFD. It is plausible that PKCδ inhibition results in a removal of GABA_A_Rs with longer decay time from the synaptic site. Further studies are warranted to further elucidate this notion.

### Activation of PKCδ activity via intracellular cascades

Unlike the effect of long-term HFD in peripheral organ that increased PKCδ transcripts and protein expression (Bezy et al., 2011), 15 days of HFD does not appear to alter PKCδ mRNA and protein expression levels in CVN^DMV^. Therefore, it seems unlikely that at this short of a time period that HFD-induced PKCδ activity in CVN^DMV^ is driven by changes in gene or protein expression. Rather, HFD may induce significant post-translational modification of PKCδ resulting in increased activation.

PKCδ is ubiquitously expressed and maintained in their inactive form in the cell. Activation of PKCδ can occur through multiple intracellular mechanisms: diacylglycerol (DAG)-dependent membrane translocation coupled with serine/threonine kinase phosphorylation and tyrosine kinase phosphorylation (Kikkawa et al., 2002). HFD exposure increases DAG levels, including in neurons (Lee et al., 2018), which in turn can drive PKC activity (Kolczynska et al., 2020). Alternatively, PKC can be activated downstream of Gq-coupled receptor in response to specific stimuli. For example, neuroactive steroids can engage PKC via Gq-coupled protein receptors signaling cascades to alter GABAergic inhibition (Lemons et al., 2025). Therefore, continued studies should investigate how intracellular signaling cascades are engaged in CVN^DMV^ to mediate the impact of HFD.

### Limitation of the study

This study used pharmacological approaches to determine PKC and PKCδ involvement in regulating tonic GABA current in CVN^DMV^. Despite widely used as a PKCδ inhibitor, rottlerin has off-target actions that may confound experimental results (Soltoff, 2007). Notably, rottlerin can activate large conductance potassium channel (BKCa) (Cordeiro et al., 2015) which could induce hyperpolarization due to potassium efflux and Ca^2+^ dependent that mimic reduction of tonic current. However, this study mitigated this secondary effect of rottlerin using Cs-gluconate internal solution which blocks potassium channels, including BKCa (Latorre et al., 2017).

### Conclusion and future directions

In summary, this study reveals the critical role of PKCδ as a regulator of tonic GABA current in CVN^DMV^ in response to metabolic challenge. Activation of this novel PKC isoform is dependent on HFD, since inhibition of PKCδ did not alter tonic current in NFD. Blocking clathrin-mediated endocytosis did not abolish the effect of PKCδ inhibition in normalizing tonic current in HFD, suggesting that PKCδ-dependent phosphorylation does not act by preventing receptor internalization, but rather through alternative mechanisms. Future studies investigating phosphorylation sites of PKCδ on GABA_A_R subunit could aid in addressing this question. How HFD drives PKCδ activity likely occurs at post-translational level since there were no changes in PKCδ mRNA and protein expression in CVN^DMV^ after 15 days of HFD. PKCδ activation and translocation are dependent on DAG, which abundance is directly influenced by HFD, and upstream Gq-protein coupled receptor cascade. Identifying the post-translational mechanisms that link HFD to PKCδ activity will provide the mechanism underpinning blunted cardiac parasympathetic output. Together, findings of this study highlights PKCδ-dependent phosphorylation of GABA_A_Rδ in CVN^DMV^ as a critical mechanism underpinning 15 days of HFD impact on cardiac vagal motor output. Future in vivo studies are warranted to investigate PKCδ as a potential therapeutic target for cardiovascular disease related to HFD.

## Authors contribution

YBW and CRB – conceptualization, YBW, VQC, CDR, MM, MJ, JNC, and CRB – data collection and analysis, YBW, MM, JNC and CRB – data interpretation, YBW – wrote the original draft of the manuscript, YBW, MM, JNC and CRB – wrote and edited the manuscript. All authors approved the final version of this manuscript.

## Acknowledgements

We thank Drs. Chiou-Miin Wang and Chun-Liang Chen at the UT Health San Antonio BASiC for their help in designing and managing the single-cell mRNA analysis.

## Methods

### Ethical approval

All procedures and experiments on animals were approved by the University of Missouri-Columbia Animal Care and Use Committee Protocol #43522. All procedures were performed according to the Guide for the Care and Use of Laboratory Animals and ARRIVE Guidelines.

### Animals

Male C57BL/6J (6-12 weeks old, #000664, Jackson Laboratory, Bar Harbor, ME, USA; RRID: IMSR_JAX:000664) were used in the experiments. Colonies are established in-house at the Dalton Cardiovascular Research Center animal facility. Mice were group-housed up to five per cage on a 12:12 light/dark cycle with ad libitum access to food and water. The number mice used is reported correspondingly in each figure legend.

### High-fat diet feeding

Animals were acclimatized to a normal fat diet (NFD; 10% kcal from fat, D12450Bi, Research Diets Inc, New Brunswick, NJ, USA) for a week prior to any experimental procedures. Animals underwent surgery to a retrogradely label cardiac vagal motor neuron, followed by 3 days of recovery period. Mice were then randomly assigned into two diet groups, a high-fat diet (HFD; 60% kcal from fat, D12492i, Research Diets Inc.) or remained on NFD and maintained for 15 days. Diet was matched based on protein content. HFD consisted of 20% kcal protein, 60% kcal fat, and 20% carbohydrate, with an energy density of 5.21 kcal/g. NFD consisted of 20% kcal protein, 10% kcal fat, and 70% carbohydrate, with an energy density of 3.82 kcal/g.

### Retrograde tracing

Mice were anesthetized with ketamine/xylazine (100/10 mg/kg, intraperitoneal). For cardiac retrograde tracing, animals were intubated using a small animal ventilator (SAR-830/AP; CWE Inc., Ardmore, PA, USA) equipped with mouse attachment (1201020). Mice were placed in a supine position on a heating pad. A small lateral incision between the second and third ribs of the right rib cages was made to expose the thoracic cavity. Retrograde tracers (40ul), tetramethylrhodamine-5-(and-6)-isothiocyanate (rhodamine, 1%, #T490, Invitrogen, Waltham, MA, USA) for patch-clamp electrophysiology, or fluorogold (FG, 1%, Fluorochrome, Denver, CO, USA) for immunohistochemistry, was injected into the pericardial fat pad near the posterior right atrioventricular junction where cardiovagal nerve endings terminate. The incision was closed using suture, and the animals were removed from the ventilator. All animals were returned to their home cage for recovery after surgery and monitored for 3 days. carprofen (10 mg/kg) and buprenorphine (0.1 mg/kg) were administered as post-operative analgesics.

### Electrophysiology

#### Preparation of acute brain slice

Mice were anesthetized with isoflurane to effect (i.e. lack of the tail-pinch reflex response) and decapitated. The brainstem was collected and 300µm thick coronal slices were prepared using a vibratome (VT1000S, Leica Biosystems, Buffalo Grove, IL, USA) in ice-cold artificial cerebrospinal fluid (ACSF) containing (in mM): 124 NaCl, 3 KCl, 26 NaHCO3, 1.4 NaH2PO4, 11 D-glucose, 1.3 CaCl2, 1.3 MgCl2 and 1 kynurenic acid (290-305 mOsm; pH 7.3-7.4). Slices were incubated in warmed (30-33°C) ACSF for 20 minutes in a holding chamber, followed by incubation at room temperature. All ACSF constituents were obtained from Sigma-Aldrich. The solution was aerated with carbogen (5% CO2/95% CO2).

#### Whole-cell patch-clamp recordings

Brain slices were transferred to a recording chamber, and neurons were visualized using Olympus BX51WI microscope (Melville, NY, USA). Whole-cell patch-clamp recordings were performed under visual control with infrared illumination and differential interference contrast (IR-DIC). Voltage-clamp configuration was used to identify GABAergic neutrotransmission inhibitory postsynaptic current (IPSC) and tonic current. A continuous perfusion of warmed aerated ACSF containing 1 mM kynurenic acid was maintained throughout recordings. Borosilicate glass pipettes (2-5 MΩ; King Precision Glass, Claremont, CA, USA) were pulled using P-97 instrument (Sutter Instruments, Novato, CA, USA) and filled with Cs-gluconate internal solution containing (in mM): 130 Cs-gluconate, 1 NaCl, 5 EGTA, 10 HEPES, 1 MgCl_2_, 1 CaCl_2_, and 2-3 Mg-ATP (pH 7.1-7.2). Cs^+^ was used as the primary cation carrier to block K^+^ currents, which allowed consistent voltage clamp at depolarized membrane potentials and diminished any influence of postsynaptic GABA_B_R during recordings. A minimum of 10-minutes equilibration period was done before recording. Patch-clamp recordings were acquired using Multiclamp 700B amplifier (Axon Instruments, Molecular Devices, Union City, CA, USA) with DigiData 1440B analog digital converter and pClamp v10.6 (Axon Instruments, Molecular Devices, Union City, CA, USA). Signals acquired at 20 kHz for voltage-clamp and low-pass filtered at 3 kHz. Both phasic and tonic inhibitory currents mediated by GABA_A_R were examined at a holding potential of 0 mV.

Glass electrodes were filled with Biocytin (0.2%, #B4261 Sigma-Aldrich, Burlington, MA, USA) was added to the internal solution to allow visualization of recorded neurons and confirmation of choline acetyltransferase (ChAT)-positive motor neurons. After recording, brainstem slices were collected and fixed in 4% paraformaldehyde (PFA) overnight, followed by 30% sucrose for two days before cryosection and immunohistochemistry.

#### Data Analysis

Analysis of recordings was done using Clampfit (Molecular Devices, San Jose, CA, USA) and Mini Analysis v6.0.3 (Synaptosoft, Decatur, CA, USA). Phasic GABA_A_R neurotransmission was determined based on analysis of 2-minutes of recording before BIC treatment. All synaptic events were used to assess IPSC frequency, but only single peak events were used to determine IPSC amplitude and decay time. IPSC amplitude and decay time analysis were performed under a requirement of a minimum number of 4 cells per group with a minimum of 30 single IPSC events per cell (IPSC frequency ≥ 0.25 Hz). Tonic current amplitudes were determined based on the difference between average holding current in 2-minutes of recording before and after BIC treatment. Tonic currents were normalized to whole-cell capacitance to correct for small differences in cell size and are presented as tonic current density (pA/pF). All recordings were discarded if series resistance was >25 MΩ or changed by >20% throughout the course of the experiment. Mean series resistance was 10.84±3.29 MΩ.

#### Drugs preparation

Drugs used in this study were obtained from Tocris Bioscience (Minneapolis, MN, USA). GABA_A_R antagonist, bicuculline methiodide (BIC; 30µM, #2503); pan-PKC inhibitor, GF 109203X (GFX; 500 nM #0741); PKCδ inhibitor, rottlerin (10 µM, #1610); clathrin-dependent endocytosis blocker, dynasore (80-100µM, #2897). All drugs were made according to the manufacturers. All drugs were applied in the perfusate for a minimum of five minutes until a steady state was reached.

### Single-cell qRT-PCR

Single-cell qRT-PCR was performed by the UTHSA Bioanalytic and Single-Cell Core (BASiC) as described previously (Wang et al., 2025). Briefly, collected cells were thawed and lysed at 75°C for 10 min, followed by DNase I treatment to remove genomic DNA contamination. RT-PCR was then performed using the CellsDirect™ One-Step qRT-PCR Kit (#11753-100, Invitrogen) on a BioMark HD MX/HX 12×12 chip microfluidics system (Fluidigm Inc., BMKHDPKG-MH; South San Francisco, CA). Primers for PKCδ were designed by UTHSA BASiC. Target genes were amplified on the BioMark HD MX/HX system using 1X SsoFast EvaGreen supermix with low ROX (Bio-Rad, N172-5211; Hercules, CA, USA) and 1X DNA binding dye sample loading reagent (Fluidigm, PN 100-3738). GAPDH and β-actin were selected as housekeeping genes. Universal mouse RNA (200 pg; BioChain, R4334566-1; Newark, CA, USA) served as a positive control. Human embryonic kidney (HEK) cells and no-template reactions served as negative controls. PCR amplification was detected on a Fluidigm BioMark HD with a FlexSix IFC using Delta Gene assays (Fluidigm PN 100-7717 B1). Detection was limited to 40 cycles. Targets that did not reach threshold within this range were considered non-detectable.

### Endocytosis Assay

Acute brainstem slices were collected following procedure for electrophysiological recording. Brainstem slices from NFD and HFD groups were incubated in ACSF containing THIP (3 µM), with vehicle (0.05% DMSO) or rottlerin (10 uM) for 2 hours, followed by pHrodo green dextran 10,000 MW (20 uM, P35368, Invitrogen, Waltham, MA, USA) for 2.25 hours. Slices were aerated with carbogen (5% CO_2_/95% CO_2_) and maintained at 30-33°C. After incubation, slices were collected and fixed in 4% PFA in 0.01 M phosphate-buffered saline (PBS) in for two days, followed by cryoprotection in 30% sucrose in PBS for two days at 4°C, prior to cryosection and immunohistochemistry.

### RNAscope

One week after pericardial Fluorogold injections, animals were deeply anesthetized with ketamine (20 mg/kg) and xylazine (2 mg/kg) diluted in sterile saline. Animals were then transcardially perfused with 0.9% saline containing heparin, followed by 10% neutral buffered formalin (NBF; Sigma-Aldrich, HT501128-4L). Brains were collected, post-fixed in 10% NBF for 24 hr at room temperature and then transferred to 30% sucrose for 48 hr at 4 °C on a shaker. Coronal brain sections (30 or 35 μm) were cut using a microtome (SM2010R, Leica Biosystems).

For RNA FISH, sections were first rinsed in PBS, mounted onto precleaned Superfrost Plus microscope slides (Fisher Scientific, catalog no. 12-550-15), and allowed to dry overnight. On the following day, an ImmEdge Hydrophobic Barrier Pen was used to outline the tissue sections. Sections were incubated with Protease IV in a HybEZ II Oven for 30 min at 40 °C and then hybridized with the RNAscope™ Probe-Mm-Prkcd (channel 1, manual assay; catalog no. 441791) for 2 hr at 40 °C. Slides were subsequently incubated with AMP 1–3, HRP-C1, HRP-C2, HRP-C3, and HRP Blocker for 15–30 min at 40 °C, as previously described (Wang et al., 2012). Probe signal was visualized using FITC (PerkinElmer), while Fluorogold was detected by its native fluorescence. Images were acquired using a Zeiss LSM 800 confocal microscope.

### Transcardiac perfusion

Mice were over anaesthetized with isoflurane until the toe-pinch reflex was not present. Mice were perfused transcardially with ice-cold PBS with heparin, followed by ice-cold 4% PFA. Brains were collected and fixed in 4% PFA for at least 24hr, followed by cryoprotection in 30% sucrose, prior to cryosection and immunohistochemistry.

### Immunohistochemistry

Brains and brainstem slices were frozen embedded in Tissue-Plus O.C.T. compound (#23-730-571, Fisher HealthCare, Houston, TX, USA). Brains and brainstem slices were sectioned in serial coronal sections of 40 μM using a cryostat (Leica Biosystems, Nussloch, Germany) for free-floating immunostaining. Slices were blocked using normal donkey serum (#017-000-121, Jackson ImmunoResearch Laboratories, West Grove, PA, USA; RRID: AB_2337258), followed by incubation of primary antibody at 4°C overnight, and 4 hours of secondary antibody (1:200) at room temperature. Primary antibodies used were rabbit anti-GABA_A_Rδ (1:50; #868A-GDN, PhosphoSolutions, Denver, CO USA; RRID: AB_2631037), goat- anti choline-acetyl transferase (1:250; #AB144P, Sigma-Aldrich, St. Louis, MO, USA; RRID: AB_11212924), rabbit-anti PKCδ (1:500; #ab182126, Abcam, Waltham, MA, USA; RRID: AB_2892154). Secondary antibodies used were donkey anti-rabbit Alexa Fluor 568 (#A-10042, Invitrogen, Waltham, MA, USA; RRID: AB_2534017), donkey anti-goat Alexa Fluor 568 (#A-11057, Invitrogen, Waltham, MA, USA; RRID: AB_142581), and Alexa Fluor Plus 488 (#A-32814, Invitrogen, Waltham, MA, USA; RRID: AB_2762838). Biocytin-filled neurons were processed via Texas Red-Avidin D staining (1:400; #A-2206, Vector Laboratories, Newark, CA, USA: RRID: AB_2336751). Slices were then mounted using Vectashield antifade mounting medium without DAPI (#H-1000-10, Vector Laboratories, Newark, CA, USA; RRID: AB_2336789) for imaging using Keyence BZ-X810 (Osaka, Japan).

Image acquisition parameters, including exposure time and intensity, were identical across groups; brightness and contrast in all images were modified together and identically for figure presentation. Negative controls were run without primary antibody. Tissue from all groups were run in parallel with the same antibody cocktails. Image analyses and mean intensity quantification were performed using ImageJ v1.54p and Cell Profiler v4.2.8 (Stirling et al., 2021).

### Statistical Analyses

Statistical analyses were performed using GraphPad Prism v11.0.2. Results are reported as mean ± standard deviation (SD). Tests for statistical significance were reported accordingly in results section. Data distribution normality was determined using Shapiro-Wilk test. Parametric data with equal variance were analyzed using either an unpaired t-test or two-way ANOVA with Sidak’s multiple comparisons post hoc. Data with two groups and unequal variance were analyzed using unpaired Welch’s t-test. Non-parametric data with more than two groups were compared with a Kruskal Wallis with a Dunn’s multiple comparison post hoc. N numbers and p- values are reported in the results section accordingly. P-value ≤0.05 is considered as statistically significant.

